# Bioluminescence Imaging of Potassium Ion Using a Sensory Luciferin and an Engineered Luciferase

**DOI:** 10.1101/2024.03.13.581057

**Authors:** Shengyu Zhao, Ying Xiong, Ranganayakulu Sunnapu, Yiyu Zhang, Xiaodong Tian, Hui-wang Ai

**Author notes:** Corresponding Author Hui-wang Ai. National Cancer Institute, 1050 Boyles St, Frederick, Maryland 21702, USA. These two authors contributed equally to this work (Shengyu Zhao, Ying Xiong).

## Abstract

Bioluminescent indicators are power tools for studying dynamic biological processes. In this study, we present the generation of novel bioluminescent indicators by modifying the luciferin molecule with an analyte-binding moiety. Specifically, we have successfully developed the first bioluminescent indicator for potassium ions (K^+^), which are critical electrolytes in biological systems. Our approach involved the design and synthesis of a K^+^-binding luciferin named potassiorin. Additionally, we engineered a luciferase enzyme called BRIPO (bioluminescent red indicator for potassium) to work synergistically with potassiorin, resulting in optimized K^+^-dependent bioluminescence responses. Through extensive validation in cell lines, primary neurons, and live mice, we demonstrated the efficacy of this new tool for detecting K^+^. Our research demonstrates an innovative concept of incorporating sensory moieties into luciferins to modulate luciferase activity. This approach has great potential for developing a wide range of bioluminescent indicators, advancing bioluminescence imaging (BLI), and enabling the study of various analytes in biological systems.

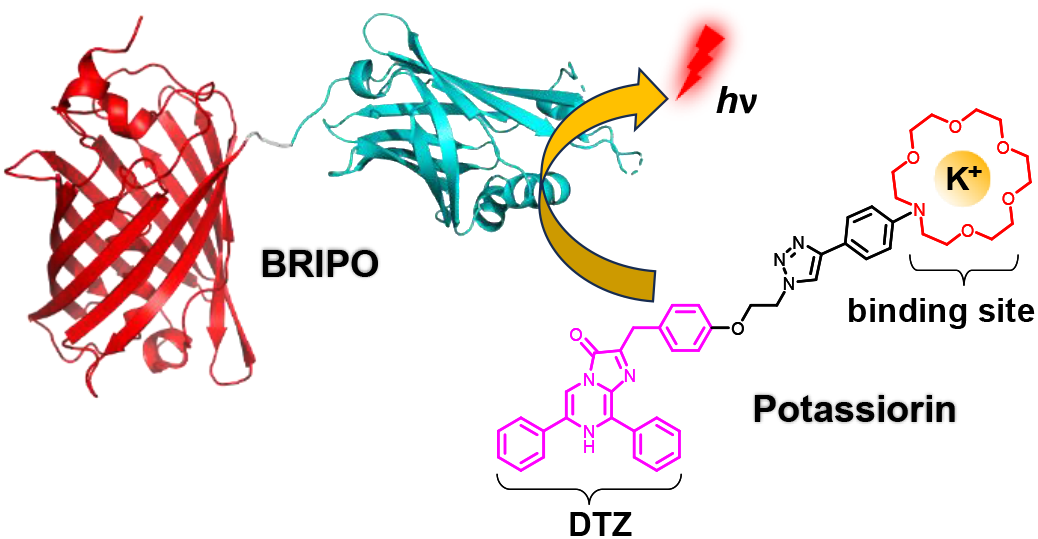

## Introduction

Fluorescent indicators have revolutionized our understanding of cellular processes by enabling real-time visualization of dynamic events in living systems.^1,2^ These indicators have become indispensable tools for researchers in various fields, facilitating significant discoveries and advancements in our understanding of life processes. Bioluminescence imaging (BLI) complements fluorescence imaging and offers distinct advantages, such as a superior signal-to-noise ratio and reduced background noise.^3-5^ BLI operates through a biochemical reaction involving the oxidation of a substrate (luciferin) by an enzyme (luciferase), allowing photon emission without external light excitation. This unique characteristic not only eliminates concerns of autofluorescence and phototoxicity but also enables deeper tissue imaging.^5^ Consequently, BLI is particularly attractive for studying biological processes in thick tissue and live animals.^6,7^ Despite its potential, the progress of BLI in visualizing biological activities is impeded by the limited availability and undesirable properties of current bioluminescent indicators.

Firefly luciferase (FLuc) is a widely used bioluminescent label. Previous research has developed a set of bioluminescent indicators by modifying the substrates of FLuc with functional groups to inhibit their bioluminescence activity.^8-11^ These modified substrates, known as caged luciferins, are designed to undergo uncaging reactions in the presence of specific molecules or enzymes, enabling the detection of biological activities (**Figure 1a**). However, these indicators rely on the availability of ATP since FLuc consumes ATP during the bioluminescence process.^4,12^ Therefore, there is a concern that they may disrupt cell physiology since ATP serves as both a vital energy currency and a crucial signaling molecule in living systems.^13^

**Figure 1.**
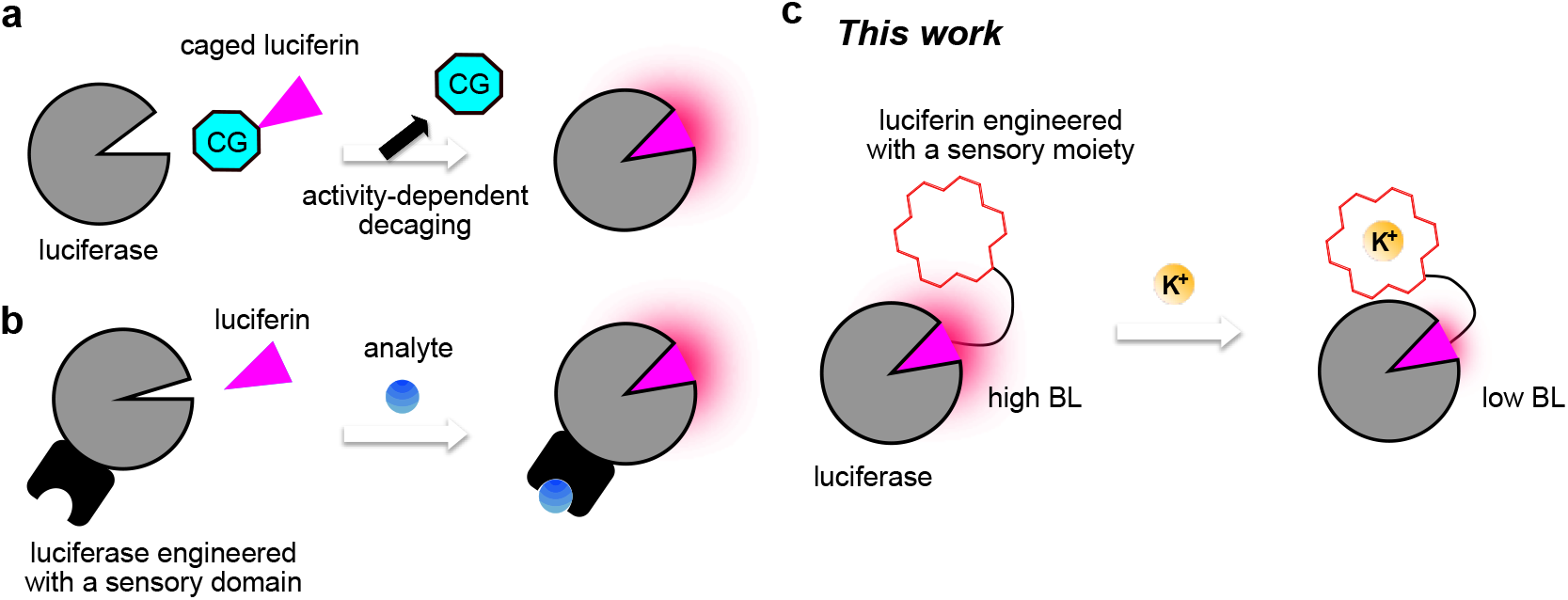
Mechanistic comparison of this work with other common bioluminescent indicators of *in vivo* imaging importance. (**a**) Reaction-based bioluminescent indicators: a caged luciferin is utilized, where a specific activity can remove the caging group and activate the luciferin, allowing for the specific detection of bioactivity. (**b**) Sensory luciferase-based bioluminescent indicators: a luciferase is engineered to be responsive to a specific analyte by strategically inserting and fusing a sensory domain to the luciferase. (**c**) Sensory luciferin-based bioluminescent indicators (this work): an analyte-binding moiety (*e*.*g*., a K^+^-binding crown ether) is strategically introduced to the luciferin, leading to the analyte-responsive modulating of the bioluminescence reaction. CG, caging group; BL, bioluminescence.

In this context, luciferases derived from marine organisms have emerged as promising candidates for indicator development.^14,15^ These luciferases utilize coelenterazine (CTZ) as their natural luciferin and do not require ATP for their activity.^4^ Among them, the NanoLuc system derived from the deep-sea shrimp *Oplophorus gracilirostris* has gained significant attention.^16^ This system offers additional advantages such as a small protein size, high enzyme stability, and a remarkable >150-fold increase in luminescence compared to traditional luciferases. In the presence of the synthetic luciferin furimazine, NanoLuc emits intense blue photons at around 450 nm. Further research has led to the development of red-shifted variants of NanoLuc, achieved through redesigned synthetic luciferins or fusion with red-emitting fluorescent proteins (FPs) for resonance energy transfer (RET).^17-21^

For indicator development, NanoLuc is frequently utilized as a RET donor. It has been fused with sensory domains and a RET acceptor to achieve RET efficiency modulation and ratiometric responses.^22-26^ However, these RET-based indicators face challenges in achieving a wide dynamic range and are better suited for *in vitro* assays rather than *in vivo* BLI applications due to the strong attenuation of NanoLuc’s blue emission by mammalian tissue.^4,5^

Another promising approach involves directly incorporating sensory domains into the structure of NanoLuc or its derived luciferases (**Figure 1b**).^21,27,28^ This results in the modulation of luciferase activity through structural changes upon analyte binding. In certain cases, the luciferases are further fused with red-emitting FPs to achieve red-shifted emission for better tissue penetration. This strategy has led to the development of several bioluminescent sensors for calcium ions (Ca^2+^) and neurotransmitters.^21,27-30^ However, this approach necessitates the identification of appropriate protein-based sensory domains for specific analytes of interest, and the engineering process can be tedious with unpredictable outcomes.

In this study, we aimed to expand the strategies for generating bioluminescent indicators. Specifically, we explored the method of modifying luciferins with sensory moieties (**Figure 1c**). We successfully designed and synthesized a luciferin called potassiorin, which selectively binds to potassium ions (K^+^), an essential electrolyte in living systems.^31^ Additionally, we engineered a luciferase named BRIPO (<u>B</u>ioluminescent <u>R</u>ed Indicator for <u>Po</u>tassium) to work in conjunction with potassiorin, producing bioluminescence signals responsive to the physiological concentrations of K^+^. To our knowledge, this development represents the first bioluminescent indicator for K^+^, expanding the capability of monitoring K^+^ in living systems. Our indicator was thoroughly tested in diverse settings, consistently demonstrating the capability in real-time monitoring of K^+^ dynamics. Furthermore, we showcased the extension of the strategy by deriving a bioluminescent indicator for sodium ions (Na^+^). Overall, our study not only presents a valuable bioluminescent indicator for studying K^+^ physiology but also introduces a powerful approach to designing bioluminescent indicators.

## RESULTS

### Design and synthesis of potassiorin

In a previous study, we introduced a NanoLuc variant called teLuc, which emits teal bioluminescence at around 500 nm when combined with the synthetic luciferin DTZ (**Figure 2a**).^18^ Notably, DTZ could be readily synthesized from commercially available reagents in just two steps with a good yield. Due to the red-shifted emission of teLuc compared to NanoLuc, we successfully developed BREP (**Figure 2b**),^28^ a fusion construct between teLuc and a red FP (RFP) mScarlet-I,^32^ emitting approximately 60% of its total emission above 600 nm. BREP enables deep-tissue photon penetration and has emerged as one of the most powerful luciferases for *in vivo* BLI. Building upon these findings, we selected DTZ and BREP as the foundations for our current study.

**Figure 2.**
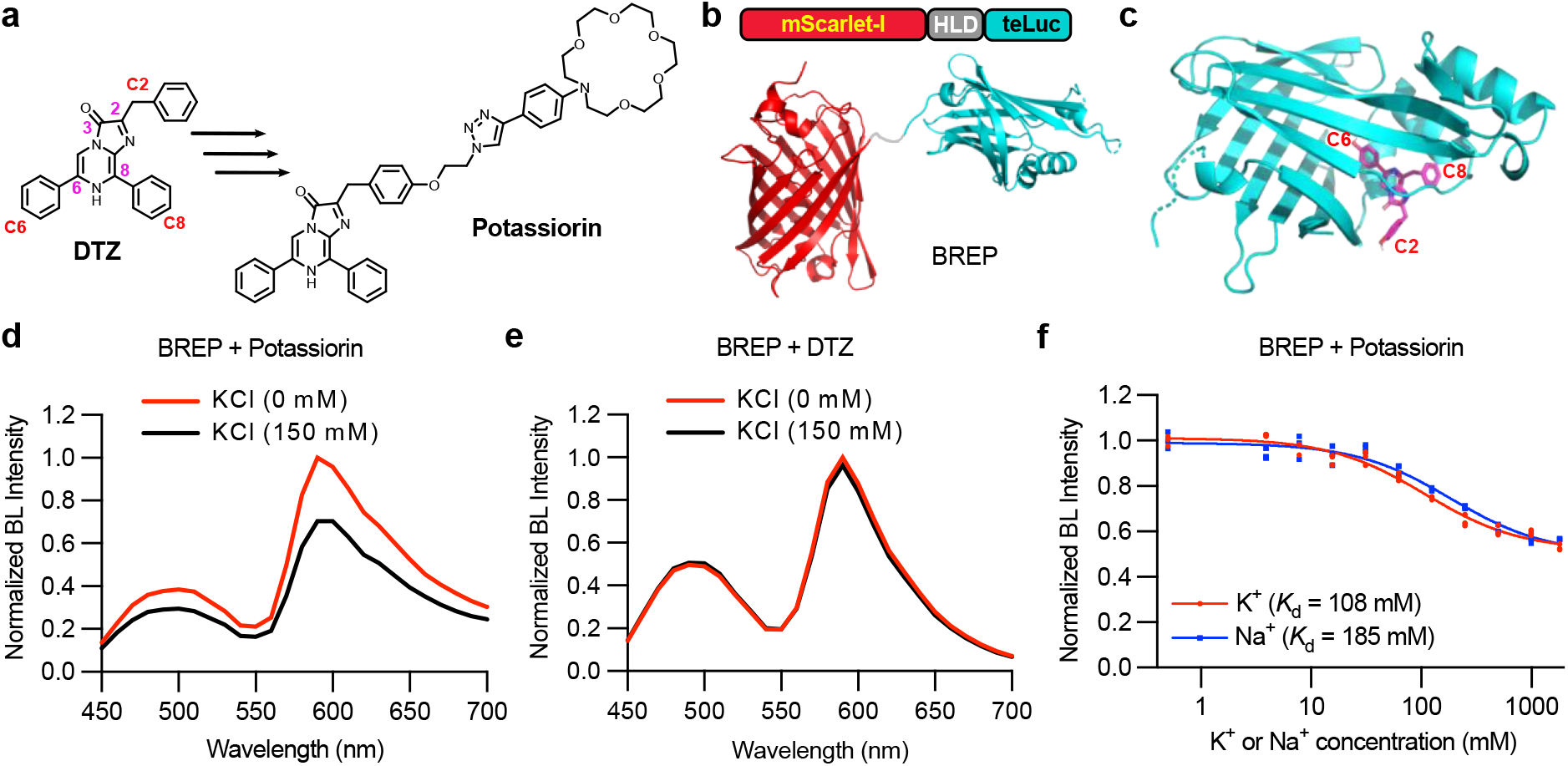
Design of potassiorin and initial evaluation with BREP luciferase. (**a**) Illustration of the DTZ structure and the installation of a K^+^-binding crown ether ring to derive potassiorin. The C2, C6, and C8 derivatizations on DTZ are highlighted. (**b**) Schematic illustration of the domain arrangements of BREP, a fusion of mScarlet-I and teLuc through a three amino acid linker. (**c**) Illustration of a modeled structure of NanoLuc (cyan ribbon) in complex with CTZ (magenta sticks). The C2, C6, and C8 derivatizations on CTZ are highlighted. (**d**,**e**) Bioluminescence emission spectra of BREP in the presence of potassiorin (**d**) or DTZ (**e**) with or without 150 mM KCl. Presented are the averages from three technical replicates. (**f**) Bioluminescence intensities of BREP and potassiorin at 590 nm in the presence of the indicated concentrations of K^+^ or Na^+^. n = 3 technical replicates. A one-site binding model was used to fit the data and derive the apparent dissociation constants (*K*_d_). BL, bioluminescence.

K^+^ is the most abundant intracellular cation, with a high concentration at 140-150 mM within cells.^31^ It plays a critical role in generating functional activity in muscle cells, neurons, and cardiac tissue.^31^ To address the limited methods available for tracking K^+^ in living systems, we aimed to develop a novel bioluminescent K^+^ indicator by incorporating a K^+^-binding moiety into DTZ. Specifically, we selected a crown ether called 1-aza-18-crown-6, known for its ability to form a complex with K^+^,^33,34^ to derivatize DTZ.

At the beginning of this project, to overcome the challenge of not having a co-crystal structure of NanoLuc and its substrate, we utilized a previously generated docking structure of CTZ in NanoLuc (**Figure 2c**).^18^ From this model, we deduced that installing the 1-aza-18-crown-6 moiety through C6 or C8 of the imidazopyrazinone core of the substrate would likely result in the complete loss of bioluminescence activity due to their buried positions and the inability of the putative substrate-binding pocket to accommodate the size of the K^+^-binding moiety. However, we identified the aromatic ring at the C2 position of imidazopyrazinone as a promising site for installing the K^+^-binding moiety, as it extends outside of the putative substrate binding pocket of the luciferase enzyme (**Figure 2c**). Notably, this rationale, which was derived from the docking model, is consistent with a recently available co-crystal structure of NanoLuc and its inactive substrate analog.^35^

We designed a DTZ analog called potassiorin, which incorporates the 1-aza-18-crown-6 moiety extended through the C2 position (**Figure 2a**). The synthesis of potassiorin involved multiple steps. Briefly, starting from commercially available 4-benzyloxybenzyl alcohol (compound **1** in **Figure S1**), we synthesized 3-(4-(2-azidoethoxy)phenyl)-1,1-diethoxypropan-2-one (**6**) in five steps with an overall yield of 9.3% (**Figure S1a**). Simultaneously, we prepared *N*-(4-ethynylphenyl)aza-18-crown-6 (**10**) in three steps from *N*-phenyldiethanolamine (**7**), with an overall yield of 32.6%. The subsequent Cu(I)-catalyzed azide–alkyne cycloaddition (CuAAC) reaction between compounds **6** and **10** produced **11** in a 77% yield (**Figure S1b**). Finally, we employed our previously established procedure to synthesize 5-diphenylpyrazin-2-amine (**12**), which was then subjected to an acid-catalyzed cyclization reaction with **11**, resulting in the final product, potassiorin, with a 10% yield (**Figure S1b**).

### Initial characterization of potassiorin with BREP

After synthesizing potassiorin, we assessed the compound using purified BREP protein. We measured the emission spectra of BREP and potassiorin in the absence and presence of 150 mM K^+^ ions (**Figure 2d**). The results indicated that the presence of 150 mM K^+^ led to a reduction in bioluminescence by approximately 30%. In contrast, the bioluminescence of BREP and DTZ (as a negative control) showed only marginal changes in response to K^+^ (**Figure 2e**). Furthermore, we examined the bioluminescence of BREP and potassiorin in the presence of various concentrations of K^+^ or Na^+^. Through our analysis, we determined the apparent affinities for K^+^ and Na^+^ (i.e., the concentrations to cause 50% of the overall bioluminescence change) to be 108 mM and 185 mM, respectively (**Figure 2f**). These findings suggest that potassiorin exhibits K^+^-dependent bioluminescence, although further improvements are necessary to enhance the dynamic range and selectivity towards K^+^ over Na^+^.

### Engineering BREP into BRIPO for enhanced responsiveness

To enhance the dynamic range and selectivity, we employed random mutagenesis on BREP using error-prone PCRs. We screened the resulting gene library and selected clones exhibiting large K^+^-dependent bioluminescence changes. These clones were subjected to counter-selection to ensure minimal bioluminescence changes in response to Na^+^. We conducted seven rounds of random mutagenesis and screening but observed only marginal improvement (**Figure S2**). To further enhance performance, we turned to multi-site-directed mutagenesis. By utilizing the docking model, we identified and simultaneously mutated three amino acids (W233, S260, and V261) near the C2 position of the luciferin. Through screening this library, we successfully identified a mutant that exhibited a remarkably improved response magnitude and selectivity. This mutant, named BRIPO, contains a total of six mutations from the original BREP sequence (**Figure 3a** and **Figure S3**). Notably, the S260R and V261W mutations obtained during the final step of engineering played a crucial role in enhancing the performance (**Figure S2a**). The co-crystal structure of NanoLuc and its substrate analog reaffirmed that these mutations are close to the C2 position of the luciferin substrate (**Figure S2b**), suggesting their ability to influence the interactions between potassiorin and the enzyme.

**Figure 3.**
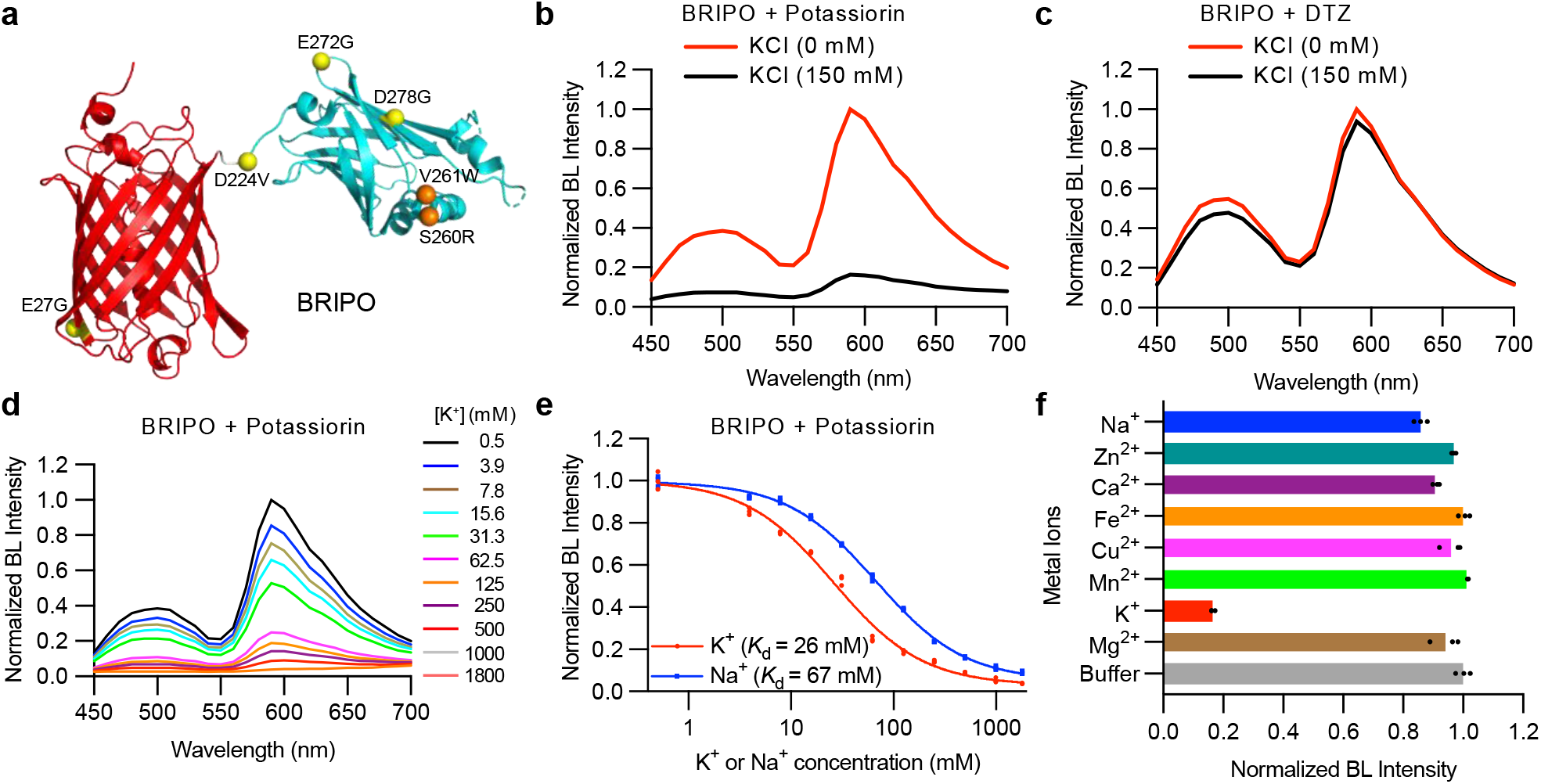
*In vitro* characterization of BRIPO. (**a**) Schematic illustration of BRIPO with mutations from BREP highlighted. (**b**,**c**) Bioluminescence emission spectra of BRIPO in the presence of potassiorin (**b**) or DTZ (**c**) with or without 150 mM KCl. Presented are the averages from three technical replicates. (**d**) Bioluminescence spectra of BRIPO and potassiorin with the indicated concentrations of KCl. Presented are the averages from three technical replicates. (**e**) Bioluminescence intensities of BRIPO and potassiorin at 590 nm in the presence of the indicated concentrations of K^+^ or Na^+^. n = 3 technical replicates. A one-site binding model was used to fit the data and derive the apparent dissociation constants (*K*_d_). (**f**) Normalized bioluminescence intensity of BRIPO and potassiorin in the presence of different metal ions: Na^+^ (15 mM), Zn^2+^ (10 μM), Ca^2+^ (2 mM), Fe^2+^ (10 μM), Cu^2+^ (100 nM), Mn^2+^ (10 μM), K^+^ (150 mM), Mg^2+^ (2 mM). n = 3 technical replicates. BL, bioluminescence.

In the presence of BRIPO and potassiorin, 150 mM K^+^ resulted in a remarkably six-fold decrease in bioluminescence intensity (**Figure 3b**). Conversely, when BRIPO was combined with DTZ, the responsiveness to K^+^ was minimal (**Figure 3c**). Further titration experiments involving varying concentrations of K^+^ and Na^+^ revealed that the apparent affinities of the BRIPO and potassiorin pair for K^+^ and Na^+^ have been altered to 26 mM and 67 mM, respectively (**Figure 3de**). Since the intracellular concentration of Na^+^ is approximately ten times lower than that of K^+^,^31^ the BRIPO-potassiorin system should exhibit sufficient specificity to accurately sense intracellular K^+^ levels over Na^+^.

To validate the specificity of the BRIPO-potassiorin system, we conducted additional tests using a range of cations at physiologically relevant or higher concentrations (**Figure 3f**). The results confirmed the system’s selectivity towards K^+^. Notably, high concentrations of Cu^2+^ led to bioluminescence quenching (**Figure S4a**), which is consistent with a previously documented mechanism involving the Cu^2+^-mediated oxidation of the luciferin.^36^ However, these high Cu^2+^ concentrations are unlikely to be physiologically relevant.^37^

Moreover, we conducted additional investigations into the potassiorin concentration dependency of BRIPO bioluminescence in the presence or absence of 150 mM K^+^ (**Figure S4b**). Our findings indicate that the presence of K^+^ enhances the binding of potassiorin to the enzyme, resulting in a lower Michaelis constant (*K*_M_). However, this increased affinity does not lead to heightened bioluminescence. Instead, we observed a reduction of nearly five-fold in the maximal photon production rate, supporting a K^+^-dependent decrease in catalysis and/or bioluminescence quantum yield.

### Imaging intracellular K^+^ dynamics in cultured cell lines

To investigate the ability of BRIPO and potassiorin to visualize K^+^ dynamics in live mammalian cells, we expressed BRIPO in HEK 293T cells and imaged the cells in a low K^+^ buffer supplemented with potassiorin. Inducing K^+^ efflux with a combination of nigericin (a K^+^ ionophore), bumetanide (an inhibitor of Na^+^/ K^+^/2Cl^−^ cotransporter), and ouabain (an inhibitor of Na^+^, K^+^-ATPase pump)^38^ resulted in an approximately 30% increase in bioluminescence (**Figure 4a-c** and **Supplementary Movie 1**). In addition, we expressed BRIPO in a HEK 293T cell line stably expressing a mouse leak K^+^ channel (mTrek) and a few other ion channels.^39^ We next used arachidonic acid to stimulate the mTrek channel.^40^ allowing K^+^ efflux to the extracellular low K^+^ space. This led to a nearly 50% increase in bioluminescence (**Figure 4d-f** and **Supplementary Movie 2**). For both cases, control experiments with DTZ did not show much change in bioluminescence (**Figure 4**). These findings support the effectiveness of the BRIPO-potassiorin system for selectively monitoring K^+^ dynamics in live mammalian cells.

**Figure 4.**
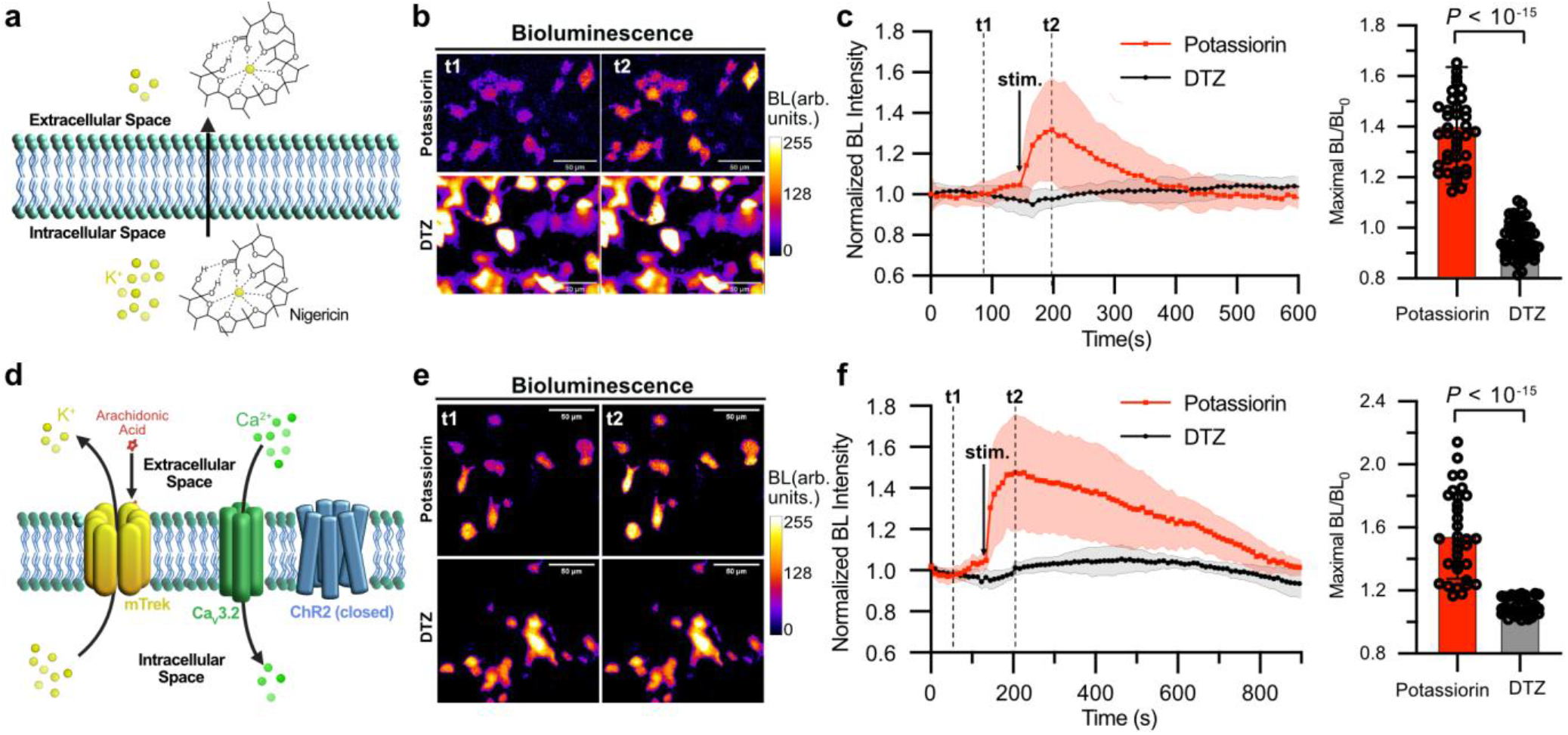
Imaging K^+^ efflux in cultured cell lines. (**a**) Schematic illustration of nigericin-mediated K^+^ efflux in HEK 293T cells in a low K^+^ buffer. (**b**) Representative pseudocolored bioluminescence images of BRIPO-expressing HEK 293T cells in the presence of potassiorin (top) or DTZ (bottom) before (left) and after (right) treatment with a combination of nigericin, ouabain, and bumetanide. Scale bar: 50 µm. (**c**) Quantification of bioluminescence intensity changes of individual cells from experiments in panel b. Data are presented as mean ± s.d. (n = 39 cells for the potassiorin group, n = 57 cells for the DTZ group). (**d**) Schematic illustration of a stable HEK 293T cell line treated with arachidonic acid to open the mTrek channel and induce K^+^ efflux. (**e**) Representative pseudocolored bioluminescence images of BRIPO-expressing HEK 293T cells stably expressing mTrek and other ion channels in the presence of potassiorin (top) or DTZ (bottom) before (left) and after (right) treatment with arachidonic acid. Scale bar: 50 µm. (**f**) Quantification of bioluminescence intensity changes of individual cells from experiments in panel e. Data are presented as mean ± s.d. (n = 33 cells for the potassiorin group, n = 42 cells for the DTZ group). In panels c and f, the baselines were corrected using a monoexponential decay model, and the *P* value was derived from unpaired two-tailed *t*-tests. The GraphPad Prism software does not provide extract *P* values below 10^-15^. BL, bioluminescence. Arb. units, arbitrary units.

### Imaging K^+^ dynamics in primary neurons and live mice

Using the BRIPO-potassiorin system, we imaged K^+^ dynamics in primary mouse neurons. We transduced the neurons with adeno-associated viruses (AAVs) and imaged them in a low K^+^ buffer with potassiorin. Glutamate was used to activate the cells, causing membrane depolarization followed by repolarization due to K^+^ channel activation and K^+^ efflux.^41^ Around 40% of the examined neurons, which are likely to express glutamate receptors, exhibited robust bioluminescence increases (**Figure 5ab**). In contrast, almost no cells in the control experiments using DTZ showed obvious changes in bioluminescence.

**Figure 5.**
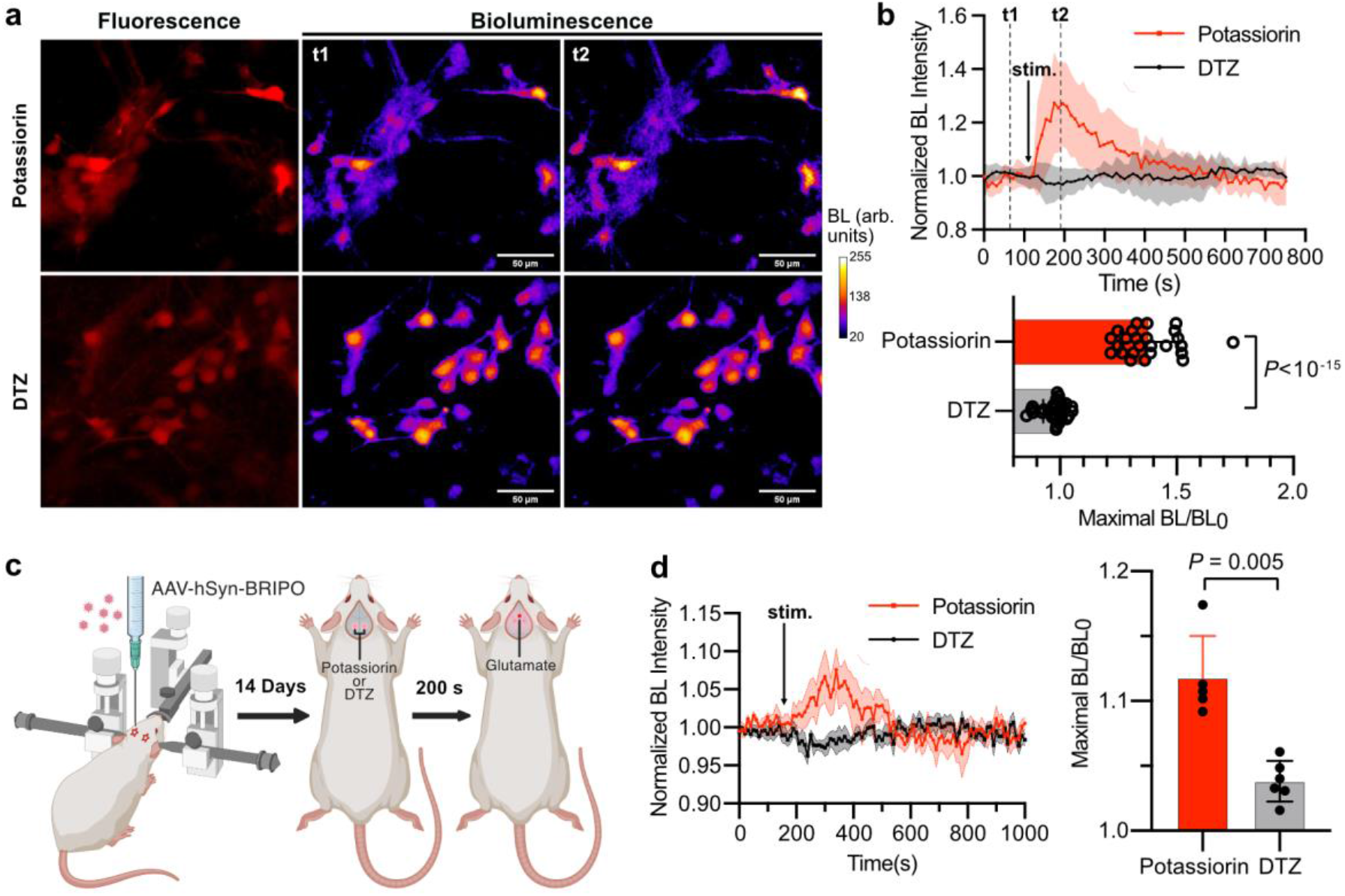
Imaging K^+^ efflux in primary mouse neurons and the brains of live mice. (**a**) Representative fluorescence and pseudocolored bioluminescence images of BRIPO-expressing primary mouse neurons. Glutamate was used to induce potassium efflux. Scale bar, 50 μm. (**b**) Quantification of bioluminescence intensity changes of individual responsive neurons upon glutamate treatment. Data are presented as mean ± s.d. (n = 25 cells for the potassiorin group, n = 31 cells for the DTZ group). (**c**) Schematic illustration of stereotactic intracranial administration of AAVs containing the BRIPO gene and other general experiment procedures. (**d**) Quantification of bioluminescence intensity changes of individual animals. Data are presented as mean ± s.e.m. (n = 5 mice for the potassiorin group, n = 6 mice for the DTZ group). In panels b and d, the baselines were corrected using a monoexponential decay model. and the *P* value was derived from unpaired two-tailed t-tests. The GraphPad Prism software does not provide extract P values below 10^-15^.

To further validate the effectiveness of the BRIPO-potassiorin system for *in vivo* imaging, we performed BLI in live mouse brains. We administered AAVs carrying the BRIPO gene into the hippocampal and cortical regions of mice. After three weeks of gene expression, we injected potassiorin and conducted time-lapse imaging on anesthetized mice placed in a dark box. Further delivery of glutamate resulted in notable increases in bioluminescence in all five mice injected with potassiorin, indicating the detection of K^+^ dynamics (**Figure 5cd** and **Supplementary Movie 3**). In contrast, control experiments using DTZ showed minimal changes in bioluminescence, which can be attributed to animal movement or alterations in blood flow. These findings provide strong evidence for the efficacy of the BRIPO-potassiorin system for *in vivo* imaging applications.

### Expanding the approach for Na^+^ sensing

To explore the potential extension of our strategy in incorporating other analyte-binding moieties into luciferins, we synthesized a potassiorin analog with a smaller crown ether ring. The size of the crown ether cavity directly affects its ability to selectively bind specific cations.^42^ In this case, we anticipated that the smaller crown ether ring would enhance the complexation with Na^+^, which is smaller than K^+^. Using a similar multi-step synthesis process as potassiorin, we successfully prepared R15-DTZ (**Figure S5**), which contains a 15-member-ring aza-crown-ether moiety. To assess the performance of R15-DTZ, we conducted assays using BREP and observed strong bioluminescence quenching induced by Na^+^ (**Figure 6a**). The BREP and R15-DTZ combination also exhibited responsiveness to K^+^, but the apparent affinity for K^+^ was approximately 6.5-fold lower than that for Na^+^ (**Figure 6b**). This suggests that the BREP and R15-DTZ pair may be well-suited for detecting extracellular Na^+^, as Na^+^ is more dominant than K^+^ in the extracellular space.^43^ Furthermore, these results indicate the potential for future modifications to develop bioluminescent indicators specifically designed for the detection of intracellular Na^+^. Overall, these findings highlight the broad applicability of our strategy in incorporating sensory moieties into luciferins, paving the way for the development of diverse bioluminescent indicators for various analytes.

**Figure 6.**
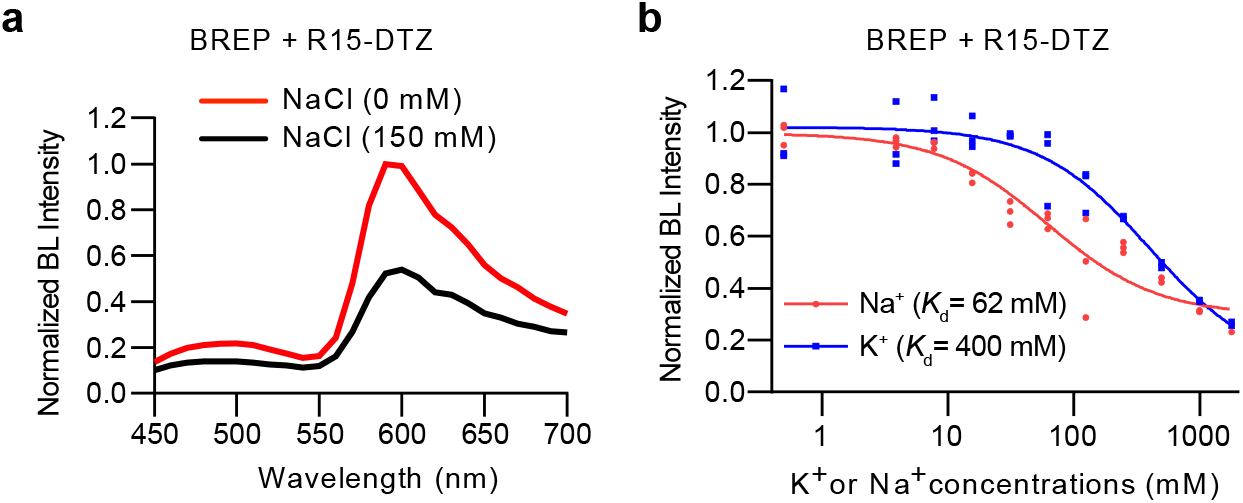
Characterization of R15-DTZ for Na^+^ sensing. (**a**) Bioluminescence emission spectra of BREP in the presence of R15-DTZ with or without 150 mM NaCl. Presented are the averages from three technical replicates. (**b**) Bioluminescence intensities of BREP and R15-DTZ at 590 nm in the presence of the indicated concentrations of K^+^ or Na^+^. n = 3 technical replicates. A one-site binding model was used to fit the data and derive the apparent dissociation constants (*K*_d_). BL, bioluminescence.

## DISCUSSION

In this study, we introduce a novel bioluminescence imaging method for studying K^+^ dynamics. We successfully developed potassiorin, a luciferin responsive to K^+^, and engineered the luciferase enzyme BRIPO to work in synergy with potassiorin. The BRIPO-potassiorin system demonstrated robust K^+^-dependent bioluminescence quenching in purified proteins, cell lines, primary neurons, and live mice, validating the effectiveness of this new system. Interestingly, the presence of K^+^ was observed to enhance the affinity of potassium for the enzyme, but it also resulted in a notable reduction in the photon production rate. This observation suggests that K^+^ binding to potassiorin allosterically modulates the enzyme activity and reduces photon production, potentially through interactions involving the mutated residues. Importantly, the K^+^-induced turn-off response provides a practical advantage: the system exhibits a bioluminescence turn-on response when cells are activated since under these conditions, K^+^ efflux occurs through K^+^ channels, leading to hyperpolarization of the cell membrane. In addition, our method overcomes the limitations of traditional K^+^-detection techniques like ion-selective electrodes,^44^ flame photometry,^45^ and fluorescent indicators,^33,34,38,41,46,47^ which face challenges in tissue- and organism-level studies. By enabling visualization and study of K^+^ dynamics from live cells to animals, our approach opens new avenues for understanding the role of K^+^ in physiology and disease.

Additionally, this study introduces a novel approach for generating bioluminescent indicators by modifying the luciferin molecule with an analyte-binding moiety. We identified that extending the luciferin molecule through the C2 aromatic ring of imidazopyrazinone is a viable option, as NanoLuc-derived luciferases display tolerance to such structural modifications while remaining sensitive to the modulation caused by analyte binding. The effectiveness of this strategy was successfully demonstrated through our development of indicators for K^+^ and Na^+^. In future studies, we plan to further enhance the sensory luciferin by incorporating ligands with higher affinity to K^+^ and Na^+^,^42,48^ aiming to improve sensitivity and enable the detection of extracellular K^+^ and intracellular Na^+^. Furthermore, we aim to expand indicator types by incorporating other sensory moieties into the luciferin structure. Overall, this research sets the stage for future advancements in bioluminescent sensors, allowing for the creation of versatile indicators that can be adapted to monitor various ions, molecules, and molecular interactions. The flexibility in sensor design opens up new avenues for broad applications BLI, which will enhance our understanding of biological systems and drive forward biomedical research.

## METHODS

### General methods and information

DNA oligos were purchased from either Integrated DNA Technologies or Eurofins Genomics. Restriction enzymes and Phusion High-Fidelity DNA polymerase were purchased from Thermo Fisher. Taq DNA polymerase was purchased from New England Biolabs. DNA sequencing was performed by Eurofins Genomics. All animal experiments were conducted following the guidelines and approval (Protocol #4196) of the University of Virginia Institutional Animal Care and Use Committee. BALB/cJ mice (#000651) were obtained from the Jackson Laboratory and housed in a temperature-controlled room (∼23°C) with a 12-hour light-dark cycle and approximately 50% humidity. At approximately six weeks of age, the mice were randomly assigned to experimental groups, ensuring a balance of both female and male animals. All ^1^H and ^13^C NMR spectra were collected on a Bruker Avance DRX 600 NMR Spectrometer at the UVA Biomolecular Magnetic Resonance Facility. Chemical shifts (δ) are given with the internal standards:^1^H (7.26 ppm) and ^13^C (77.0 ppm) for CDCl^3^; ^1^H (5.32) for CD_2_Cl_2_, and ^13^C (49.00 ppm) for CD_3_OD. Splitting patterns are reported as s (singlet), d (doublet), t (triplet), dd (doublet of doublets), and m (multiplet). Coupling constants (J) are reported in Hz. Synthetic schemes and compound numbering information are shown in **Figs. S1 & S5a**. NMR spectra for key compounds are presented in **Figs. S6 & S7**. ChatGPT was utilized to paraphrase sentences and correct grammatical errors in this manuscript.

### Synthesis of 3-(4-(2-azidoethoxy)phenyl)-1,1-diethoxypropan-2-one (6)

First, compound **2** was synthesized as a white powder from 4-benzyloxybenzyl alcohol (**1**) following a published procedure.^49^ Next, **4** was synthesized from **2** in two steps using a previous procedure.^50^ Subsequently, **6** was obtained as a colorless oil from **4** in two steps as previously reported.^51^

### Synthesis of *N*-(4-ethynylphenyl)aza-18-crown-6 (10)

Compound **10** was synthesized from *N*-phenyl diethanolamine (**7**) through several steps. First, **8** was obtained as a white solid according to the literature.^52^ Then, **9** was prepared from **8** using a published procedure.^53^ In the next step, compound **10**, which was reported previously,^54^ was obtained from **9** using a revised procedure. Briefly, **9** (600 mg, 1.63 mmol, 1 equiv.) was dissolved in 10 mL dry methanol and stirred with dry K_2_CO_3_ (899 mg, 6.52 mmol, 4 equiv.) in a 100 mL round bottom flask at room temperature. Then, 588 uL of dimethyl (1-diazo-2-oxopropyl) phosphonate (753 mg, 3.9 mmol, 2.4 equiv.) was added. The reaction mixture was stirred overnight at room temperature and monitored by thin-layer chromatography (TLC). After the reaction neared completion, 50 mL ddH_2_O was added to the reaction mixture. Then, the resulting mixture was subjected to three extractions with 50 mL of ethyl acetate each time. The organic layers were combined, dried over anhydrous Na_2_SO_4_, filtered, and concentrated under vacuum. The resulting residue was purified by silica column chromatography using an elution solvent mixture of ethyl acetate and hexane (3:10, gradually shifting to pure ethyl acetate). This yielded compound **10** as a white solid (473 mg, 1.3 mmol, 80% yield).

### Synthesis of 3-(4-(2-(4-(4-(1,4,7,10,13-pentaoxa-16-azacyclooctadecan-16-yl)phenyl)-1H-1,2,3-triazol-1-yl)ethoxy)phenyl)-1,1-diethoxypropan-2-one (11)

Compound **10** (180 mg, 0.585 mmol) and compound **6** (236 mg, 0.585 mmol) were suspended in a mixture of ddH_2_O (4 mL) and *tert*-butyl alcohol (4 mL). Then, sodium ascorbate (12.8 mg, 0.0585 mmol, freshly prepared as a 5 mL solution in ddH_2_O) was added, followed by copper (II) sulfate pentahydrate (1 mg, 0.00585 mmol, pre-dissolved in 5 mL ddH_2_O). The resulting heterogeneous mixture was vigorously stirred overnight until it cleared, and TLC analysis confirmed the complete consumption of the reactants. The reaction mixture was subsequently diluted with 20 mL of ddH_2_O and extracted three times with 20 mL of ethyl acetate. The organic layers were combined, washed with 20 mL of brine, dried over Na_2_SO_4_, filtered, and concentrated under vacuum. The resulting residue was purified by silica column chromatography using an elution solvent mixture of ethyl acetate and hexane (3:10, gradually shifting to pure ethyl acetate). This yielded 300 mg (77% yield) of pure product as a sticky light-yellow oil. ^1^H NMR-(600 MHz, CDCl_3_): δ 7.78 (s, 1H), 7.62 (d, J = 8.8 Hz, 2H), 7.10 (d, J = 8.7 Hz, 2H), 6.82 – 6.81 (m, 2H), 6.70 ( d, J = 8.9 Hz, 2H), 4.74 – 4.72 (m, 2H), 4.59 (s, 1H), 4.34 – 4.32 (m, 2H), 3.80 (s, 2H), 3.71 – 3.63 (m, 26H), 3.55 – 3.50 (m, 2H), 1.21 (t, J = 14 Hz, 6H).^13^C NMR (151 MHz, CDCl_3_): δ 203.3, 156.8, 148.2, 147.8, 130.9, 126.9, 119.1, 114.6, 111.8 102.2, 70.7, 70.65, 70.62, 70.60, 68.6, 66.5, 63.3, 51.2, 49.6, 42.6, 15.0. HR-MS (C_35_H_50_N_4_O_9_): [M+H]^+^, calcd: 671.3651, found: 671.3653.

### Synthesis of potassiorin (13)

3,5-diphenylpyrazin-2-amine (**12**) was prepared from commercially available 3,5-dibromopyrazin-2-amine according to a previously described procedure.^18^ Next, a solution of compound **12** (25 mg, 0.1 mmol, 1 equiv.) and compound **11** (134 mg, 0.2 mmol, 2 equiv.) in 5 mL degassed 1,4-dioxane was prepared. Then, 0.5 mL of 6 N HCl (30 equiv.) was added to the solution. The resulting mixture was stirred at 80 °C in a sealed pressure tube (MilliporeSigma, Cat. # Z568767) for 12 hours. Afterward, the reaction was cooled down to room temperature, and the solvent was removed under vacuum. The residue was dissolved in a 1 mL solution of methanol and water (1:1, v/v). The resulting mixture was filtered through a 0.22 μm polytetrafluoroethylene (PTFE) membrane filter and further purified with a Waters Prep 150 liquid chromatography coupled with an SQ Detector 2 mass spectrometer. An XBridge BEH Amide/Phenyl OBD Prep Column (130Å, 5 μm, 30 mm × 150 mm) was used along with a gradient elution of acetonitrile and water (1:99 to 90:10) at a flow rate of 20 mL/min. The fractions containing the desired product were combined and subjected to lyophilization, resulting in the potassiorin compound as an orange powder (8 mg, 0.01 mmol, 10% yield). ^1^H-NMR (600 MHz, CD_2_Cl_2_) δ 8.66 (d, J = 20.3 Hz, 2H), 8.05 – 8.02 (m, 4H), 7.97 (dd, J = 1.5, 7.5 Hz, 2H), 7.80 (d, J = 8.7 Hz, 2H), 7.61 – 7.59 (m, 3H), 7.47 – 7.44 (m, 2H), 7.41 – 7. 38 (m, 1H), 7.25 (d, J = 8.7, 2H), 6.91 (d, J = 8.7, 2H), 4.84 (t, J = 9.9 Hz, 2H), 4.41 (t, J = 10 Hz, 2H), 4.21 (s, 1H), 3.82 – 3.60 (m, 20H), 3.26 – 3.25 (m, 4H). ^13^C NMR (151 MHz, CD_3_OD) δ 158.7, 146.9, 146.5, 142.8, 139.0, 136.6, 135.6, 134.2, 133.6, 132.8, 131.1, 131.0, 130.8, 130.3, 130.2, 130.1, 128.7, 127.9, 127.4, 125.3, 124.6, 123.8, 116.1, 111.6, 71.5, 71.4, 71.2, 70.5, 67.6, 65.0, 59.9, 51.6, 29.5. HR-MS (C_47_H_51_N_7_O_7_): [M+H]^+^, calcd: 826.3923, found: 826.3892.

### Synthesis of R15-DTZ (18)

Compound **18** was synthesized following a procedure similar to that of potassiorin (**13**). However, instead of using compound **8**, which was prepared from compound **7**, we utilized commercially available compound **14** as the starting material. The synthetic scheme is provided in **Figure S5**. In the final step, the product was obtained as an orange powder.

### Library construction and screening

To create libraries with random mutations, the BREP gene was amplified from our previously described pcDNA3-BREP plasmid (Addgene, Cat. # 172337)^28^ using Taq DNA polymerase under a previously established error-prone condition.^55^ The resulting mutated genes were then subcloned into a pBAD/His B plasmid using Gibson assembly.^56^ *E. coli* DH10B competent cells were transformed by electroporation and plated on 2xYT agar supplemented with 100 μg/mL ampicillin and 0.2% (w/v) L-arabinose. After overnight incubation at 37°C, approximately 200 μL of 25 μM potassiorin was sprayed onto the colonies on each plate. BLI was performed using a UVP BioSpectrum dark box, a Computar Motorized ZOOM lens (M6Z1212MP3), and a Teledyne Photometrics Evolve 16 EMCCD camera. Colonies displaying strong bioluminescence were selected and cultured individually in wells of 96-well plates containing 1 mL of 2xYT media supplemented with 100 μg/mL ampicillin and 0.2% (w/v) L-arabinose. After shaking at 37°C for 20 hours, bacterial cells were pelleted by centrifugation and lysed using 500 μL of Thermo Fisher Bacterial Protein Extraction Reagent (B-PER). In the initial screening stage, the bioluminescence of *E. coli* lysates was measured in the presence of potassiorin under two KCl concentrations: 0 mM and 150 mM. For this, 30 μL of each cell lysate was diluted with 50 μL of MOPS buffer (10 mM, pH 7.4) containing either 0 mM or 240 mM KCl, resulting in final KCl concentrations of 0 mM and 150 mM, respectively. Meanwhile, potassiorin (5 mM) dissolved in a premade stock solution (ethanol:1,2-propanediol = 1:1 (v/v), supplemented with 0.88 mg/mL L-ascorbic acid) was diluted to 100 μM using the MOPS buffer containing no KCl or 150 mM KCl. A 20 μL aliquot of the potassiorin solution was dispensed into each well of a microplate using an automated dispenser on a CLARIOstar Microplate Reader (BMG Labtech). After a 1-second shake, the bioluminescence spectra ranging from 450 nm to 700 nm were recorded using the plate reader equipped with a red-sensitive PMT. Mutants that exhibited extreme K^+^-dependent bioluminescence changes were chosen for subsequent screening, which focused on their resistance to Na^+^. A similar procedure as described above was employed to test the bioluminescence responses of the mutants to 30 mM NaCl versus no NaCl. The mutant showing the highest response to K^+^ and the lowest response to Na^+^ was chosen as the template for the next screening round. To construct the focused library targeting residues 233, 260, and 261, oligos containing NNK degenerate codons (where N = A, T, G, or C and K = G or T) were utilized to amplify three short gene fragments. Subsequently, Gibson assembly^56^ was employed to fuse these fragments with the predigested pBAD/His B plasmid. The remaining steps involved in library screening were identical to the procedures described above.

### Protein purification and *in vitro* assays

Recombinant proteins BREP and BRIPO were expressed and purified following a previous procedure,^28^ and the purity was verified using SDS-PAGE (**Fig. S8**). The purified proteins were diluted in MOPS buffer to a final concentration of 200 nM, with either 0 mM or 300 mM KCl. For the initial assay, 50 μL of each protein dilution was added to the wells of a 96-well plate. Then, 50 μL of potassiorin (50 μM) in MOPS buffer was dispensed into each well, resulting in final KCl concentrations of 0 mM or 150 mM. The bioluminescence spectra were recorded using the CLARIOstar Microplate Reader. For the K^+^ or Na^+^ concentration dependence assays, 50 μL of MOPS buffer containing 200 nM purified proteins and a specific concentration of NaCl or KCl was added to the wells of a 96-well plate. Then, 50 μL of potassiorin (50 μM) in MOPS buffer was injected into each well, establishing final ion concentrations ranging from 1800 mM to 0.5 mM. The bioluminescence spectra were recorded and the intensity values at 590 nm were plotted against ion concentrations. Data was fit using the one-site binding model in GraphPad Prism 9. For the ion selectivity assays, 50 μL of MOPS buffer containing 200 nM purified BRIPO and a specific metal ion was added to the wells of a 96-well plate. Then, 50 μL of potassiorin (50 μM) in MOPS buffer was injected into each well. Bioluminescence was recorded, with metal ions supplied as follows: NaCl (15 mM), ZnCl_2_ (10 μM), CaCl_2_ (2 mM), KCl (150 mM), MgCl_2_ (2 mM), MnCl_2_ (10 μM), FeCl_2_ (10 μM), CuCl_2_ (100 nM). Two additional CuCl_2_ concentrations (1 μM and 10 μM) were tested in the presence of either potassiorin or DTZ. For the substrate concentration dependence assays, 50 μL of MOPS buffer containing 200 nM purified BRIPO and either no or 300 mM KCl was added to the wells of a 96-well plate. Various volumes of MOPS buffer and potassiorin solutions were injected into each well to achieve potassiorin concentrations ranging from 25 μM to 0.5 μM. After a 1-second shake, the bioluminescence of each well was recorded. The data was fit using the Michaelis-Menten nonlinear regression function in GraphPad Prism 9.

### Characterization in mammalian cell lines

The BRIPO gene was amplified from the pBAD plasmid and inserted into a pcDNA3 vector, resulting in pcDNA3-BRIPO. HEK 293T cells (ATCC, Cat. # CRL-3216) were cultured in DMEM supplemented with 10% fetal bovine serum (FBS). A HEK 293T cell line stably expressing mTrek1 and the α1H subunit of Ca_V_3.2, provided by Dr. Paula Barrett (University of Virginia), was cultured in DMEM supplemented with 10% FBS, 1% penicillin/streptomycin, 0.4 μg/mL puromycin, and 400 μg/mL G418. The generation and characterization of this cell line were previously described.^39^ Both types of cells were transfected using a previously described procedure.^28^ Imaging was conducted 2 to 3 days later. On the imaging day, cells were rinsed three times with a lab-made cell imaging buffer (15 mM D-glucose, 0.1 mM sodium pyruvate, 0.49 mM MgCl_2_, 2 mM CaCl_2_, 0.4 mM MgSO_4_, 0.44 mM KH_2_PO_4_, 5.3 mM KCl, 4.2 mM NaHCO_3_, 0.34 mM Na_2_HPO_4_, 138 mM NaCl, 10 mM HEPES, pH 7.2). Cells were then maintained in this buffer supplemented with 50 µM potassiorin or DTZ for bioluminescence. Time-lapse imaging was performed using an inverted Leica DMi8 microscope equipped with a Photometrics Prime 95B Scientific CMOS camera. The imaging settings included a 40× oil immersion objective lens (NA 1.2), no filter cube, 2×2 camera binning, 10 s exposure with no interval, camera sensor temperature of –20 °C, and 12-bit high-sensitivity mode. For HEK 293T cells, nigericin (10 mM), bumetanide (10 mM), and ouabain (10 mM) ethanol stocks were diluted in the imaging buffer mentioned above to achieve final concentrations of 20 μM, 10 μM, and 10 μM, respectively. For the stable HEK 293T cells, arachidonic acid (10 mM) ethanol stock was diluted in the same imaging buffer to a final concentration of 20 μM. Acquired images were processed using the Fiji version of ImageJ 1.53e as described.^28^ Data were plotted, and statistical analysis was performed using GraphPad Prism 9. The baselines caused by substrate decay were corrected according to the previously described procedure.^28^

### Viral preparation and characterization in primary mouse neurons

The BRIPO gene was amplified from the corresponding pcDNA3 plasmid and subsequently inserted into a pAAV-hSyn vector, resulting in the creation of pAAV-hSyn-BRIPO. AAVs carrying the BRIPO gene were prepared using our previously reported procedure.^28^ The obtained AAV titers were about 1×10^13^ GC/mL. Following preparation, the AAVs were aliquoted and stored at –80 °C for long-term preservation. Primary mouse neurons were prepared as described.^28^ Neurons were seeded on 35 mm glass-bottom dishes coated with poly-D-lysine, supplemented with 2 mL NbActiv4 medium (BrainBits). The culture was maintained at 37 °C with 5% CO_2_. On the fourth day post-plating, half of the medium was changed to fresh NbActiv4. On the same day, 3 µL of the BRIPO virus and 1 μL of 1 M HEPES (pH 7.4) were added to each 35 mm culture dish. Neurons were imaged four days post-transduction. The growth medium was carefully replaced with 0.5 mL of the cell imaging buffer supplemented with 100 µM of potassiorin or DTZ before imaging. Time-lapse imaging was performed under the same settings described in the mammalian cell imaging section. During time-lapse imaging, glutamate dissolved in the above-mentioned imaging buffer was added to the dish at a final concentration of 1 mM. Image processing and data analysis were identical to the procedure described in the mammalian cell imaging section.

### Imaging of K^+^ dynamics in live mice

For each BALB/cJ mouse, 500 nL of AAV was delivered to both sides of the hippocampus (AP –1.7, ML ± 1.2, DV –1.5) and cortex (AP –1.7, ML ± 1.2, DV –0.5) via intracranial stereotactic injection at a flow rate of 100 nL/min. The needle remained in the brain for additional 5 min after the infusion was complete, and the wound was sealed with surgical adhesive. Two to three weeks after the virus injection, potassiorin (5 mM) or DTZ (15 mM) pre-dissolved in a stock solution (ethanol:1,2-propanediol = 1:1 (v/v), supplemented with 0.88 mg/mL L-ascorbic acid) was diluted in saline to a concentration of 25 μM. The mice were anesthetized, and 500 nL of the diluted compound was injected into the virus infusion sites. Time-lapse imaging was then performed using a UVP BioSpectrum dark box, a Computer Motorized ZOOM lens (M6Z1212MP3), and a Teledyne Photometrics Evolve 16 camera. The instrumental settings were as follows: camera sensor gain of 3, PMT gain of 600, 2×2 binning, camera sensor temperature of –20 °C, and 10 s exposure time with no interval. The ZOOM lens was set to be 100% open, 0% zoom, and 0% focus. The mice were positioned 20 cm away from the front of the lens without an emission filter. During the time-lapse imaging, the mice were briefly removed from the dark box and intracranially injected with 500 nL of glutamate (10 mM in saline) into the middle of the virus injection sites (AP –0.7, ML 0, DV –1.0) at a flow rate of 250 nL/min. The mice were immediately placed back in the dark box for subsequent imaging. Data analysis followed the same procedure described in the mammalian cell imaging section.

## ASSOCIATED CONTENT

### Supporting Information

The Supporting Information is available free of charge on the ACS Publications website.

Synthesis and characterization data, BRIPO engineering, key mutations and sequence alignment, and additional in vitro characterization. (PDF)

Movie S1. BLI of BRIPO-expressing HEK 293T cells in response to a combination of nigericin, ouabain, and bumetanide. (AVI)

Movie S2. BLI of BRIPO-expressing HEK 293T cells stably expressing mTrek and several other ion channels in response to arachidonic acid. (AVI)

Movie S3. BLI of a mouse with BRIPO expression in the brain in response to local glutamate stimulation. (AVI)

## AUTHOR INFORMATION

## Author Contributions

H.A. conceived the project. S.Z. conducted protein engineering and enzyme assays, viral preparation, animal injection, and in vitro and in vivo imaging experiments. Y.X. established the initial synthetic routes, synthesized potassiorin and R15-DTZ, and assisted S.Z. with protein engineering, enzyme assays, and cell imaging. R.S. revised the synthetic routes and synthesized potassiorin and R15-DTZ. X.T. provided DTZ. Y.Z. prepared the primary neurons, and Y.Z. and X.T. assisted S.Z. during the live animal imaging experiments. Data analysis and figure preparation were conducted by S.Z., Y.X., and R.S. The figures were revised by H.A. The manuscript was written by H.A., S.Z., Y.X., and R.S.

## Notes

While currently there is no plan to patent BRIPO and potassiorin, HA was an inventor of a patent (US Application # 15/694238) about DTZ and teLuc awarded to the University of California. Additionally, the University of Virginia filed a patent application (US Application # 17/434351) that covers BREP, with HA and YX listed as inventors. The remaining authors declare no competing interests.

## ACKNOWLEDGMENT

We acknowledge the UVA Biomolecular Magnetic Resonance Facility for technical assistance. Research reported in this publication was supported by the University of Virginia Start-up Fund and National Institutes of Health grants (R01EB035430, R01EB033172, R01DK122253, and RF1AG077773) to HA.

## ABBREVIATIONS

BRIPO: bioluminescent red indicator for potassium
BL: bioluminescence
BLI: bioluminescence imaging
FLuc: Firefly luciferase
FP: fluorescent proteins
RET: resonance energy transfer
K^+^: potassium ions
Na^+^: sodium ions
RFP: red fluorescent protein
*K*_M_: Michaelis constant
*K*_d_: apparent dissociation constant
AAVs: adeno-associated viruses.

