## Supplementary Information for "Bioluminescence Imaging of Potassium Ion Using a Sensory Luciferin and an Engineered Luciferase"

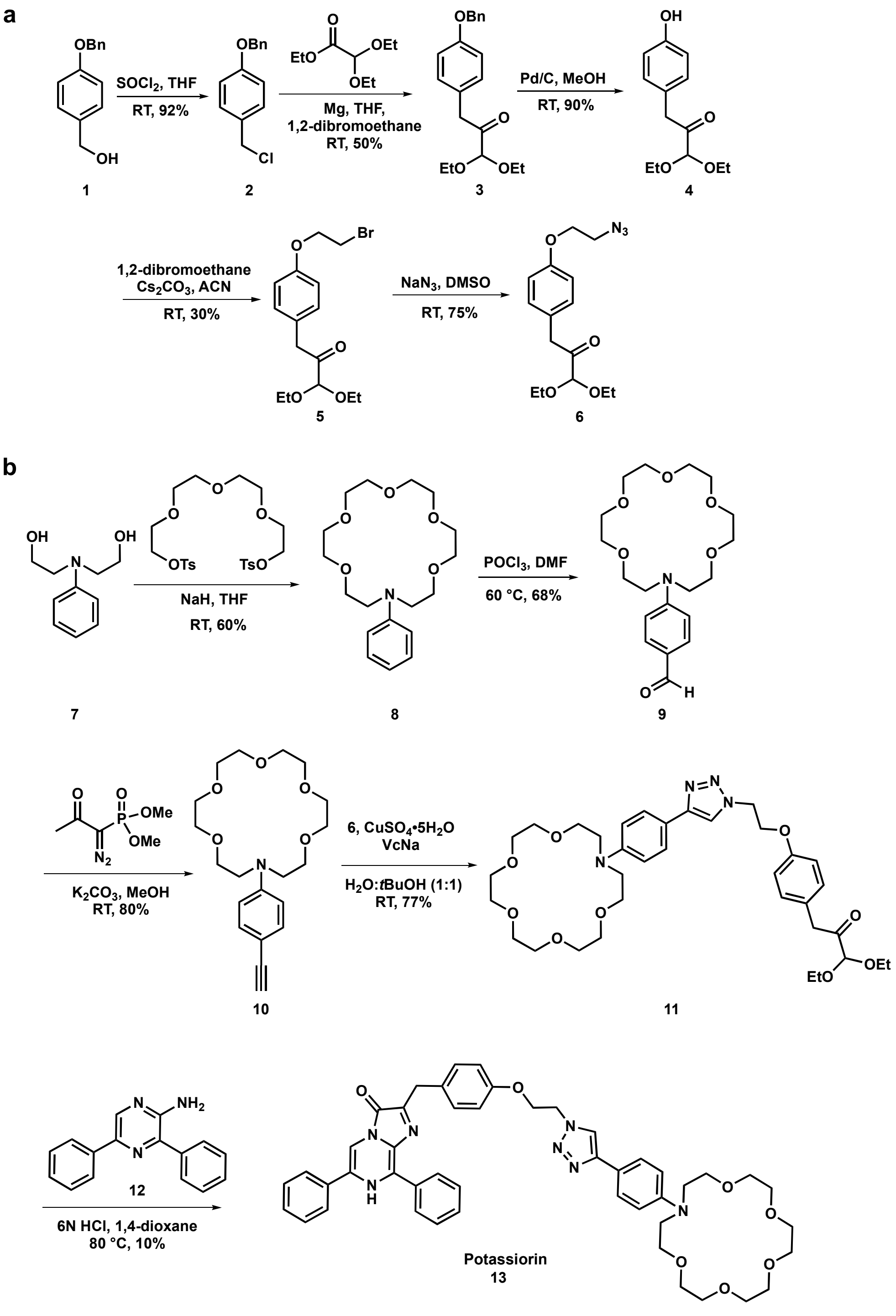


**Figure S1. Synthetic route to chemically synthesize potassiorin.** (**a**) Preparation of 3-(4-(2-azidoethoxy)phenyl)-1,1-diethoxypropan-2-one (**6**) from 4-benzyloxybenzyl alcohol (**1**). (**b**) Preparation of potassiorin from N-phenyldiethanolamine (**7**) and **6**.

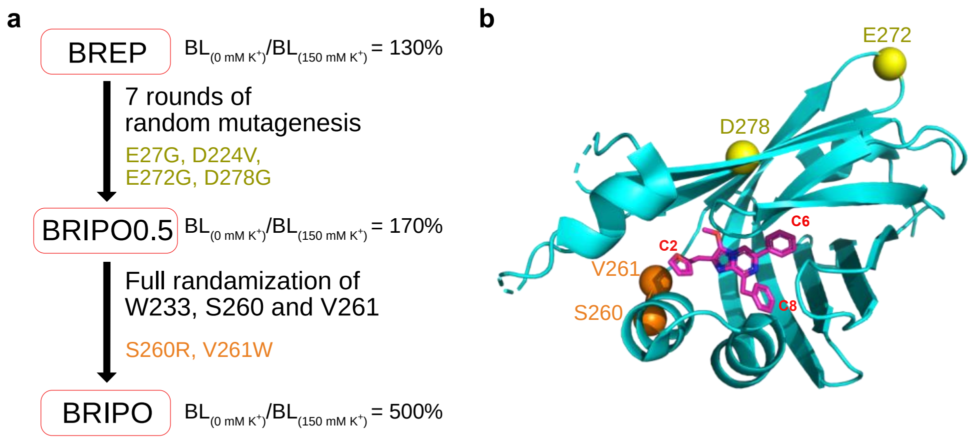


**Figure S2. Engineering of BRIPO and illustration of key mutations.** (**a**) Flowchart for the protein engineering process with mutations and dynamic ranges (determined with cell lysates) highlighted. (**b**) Structure of NanoLuc bound with an inactive 3-methoxy-furimazine luciferin analog (Protein Data Bank entry 7SNT). Residues in the luciferase domain that are mutated in BRIPO are also indicated. BL, bioluminescence.

| | 1 | 2 | 3 | 4 | 5 | 6 | 7 | 8 | 9 | 10 | 11 | 12 | 13 | 14 | 15 | 16 | 17 | 18 | 19 | 20 | 21 | 22 | 23 | 24 | 25 | 26 | 27 | 28 | 29 | 30 |
|---|---|---|---|---|---|---|---|---|---|---|---|---|---|---|---|---|---|---|---|---|---|---|---|---|---|---|---|---|---|---|
| **BREP** | M | V | S | K | G | E | A | V | I | K | E | F | M | R | F | K | V | H | M | E | G | S | M | N | G | H | E | F | E | I |
| **BRIPO0.5** | M | V | S | K | G | E | A | V | I | K | E | F | M | R | F | K | V | H | M | E | G | S | M | N | G | H | G | F | E | I |
| **BRIPO** | M | V | S | K | G | E | A | V | I | K | E | F | M | R | F | K | V | H | M | E | G | S | M | N | G | H | G | F | E | I |

| | 31 | 32 | 33 | 34 | 35 | 36 | 37 | 38 | 39 | 40 | 41 | 42 | 43 | 44 | 45 | 46 | 47 | 48 | 49 | 50 | 51 | 52 | 53 | 54 | 55 | 56 | 57 | 58 | 59 | 60 |
|---|---|---|---|---|---|---|---|---|---|---|---|---|---|---|---|---|---|---|---|---|---|---|---|---|---|---|---|---|---|---|
| **BREP** | E | G | E | G | E | G | R | P | Y | E | G | T | Q | T | A | K | L | K | V | T | K | G | G | P | L | P | F | S | W | D |
| **BRIPO0.5** | E | G | E | G | E | G | R | P | Y | E | G | T | Q | T | A | K | L | K | V | T | K | G | G | P | L | P | F | S | W | D |
| **BRIPO** | E | G | E | G | E | G | R | P | Y | E | G | T | Q | T | A | K | L | K | V | T | K | G | G | P | L | P | F | S | W | D |

| | 61 | 62 | 63 | 64 | 65 | 66 | 67 | 68 | 69 | 70 | 71 | 72 | 73 | 74 | 75 | 76 | 77 | 78 | 79 | 80 | 81 | 82 | 83 | 84 | 85 | 86 | 87 | 88 | 89 | 90 |
|---|---|---|---|---|---|---|---|---|---|---|---|---|---|---|---|---|---|---|---|---|---|---|---|---|---|---|---|---|---|---|
| **BREP** | I | L | S | P | Q | F | M | Y | G | S | R | A | F | I | K | H | P | A | D | I | P | D | Y | Y | K | Q | S | F | P | E |
| **BRIPO0.5** | I | L | S | P | Q | F | M | Y | G | S | R | A | F | I | K | H | P | A | D | I | P | D | Y | Y | K | Q | S | F | P | E |
| **BRIPO** | I | L | S | P | Q | F | M | Y | G | S | R | A | F | I | K | H | P | A | D | I | P | D | Y | Y | K | Q | S | F | P | E |

| | 91 | 92 | 93 | 94 | 95 | 96 | 97 | 98 | 99 | 100 | 101 | 102 | 103 | 104 | 105 | 106 | 107 | 108 | 109 | 110 | 111 | 112 | 113 | 114 | 115 | 116 | 117 | 118 | 119 | 120 |
|---|---|---|---|---|---|---|---|---|---|---|---|---|---|---|---|---|---|---|---|---|---|---|---|---|---|---|---|---|---|---|
| **BREP** | G | F | K | W | E | R | V | M | N | F | E | D | G | G | A | V | T | V | T | Q | D | T | S | L | E | D | G | T | L | I |
| **BRIPO0.5** | G | F | K | W | E | R | V | M | N | F | E | D | G | G | A | V | T | V | T | Q | D | T | S | L | E | D | G | T | L | I |
| **BRIPO** | G | F | K | W | E | R | V | M | N | F | E | D | G | G | A | V | T | V | T | Q | D | T | S | L | E | D | G | T | L | I |

| | 121 | 122 | 123 | 124 | 125 | 126 | 127 | 128 | 129 | 130 | 131 | 132 | 133 | 134 | 135 | 136 | 137 | 138 | 139 | 140 | 141 | 142 | 143 | 144 | 145 | 146 | 147 | 148 | 149 | 150 |
|---|---|---|---|---|---|---|---|---|---|---|---|---|---|---|---|---|---|---|---|---|---|---|---|---|---|---|---|---|---|---|
| **BREP** | Y | K | V | K | L | R | G | T | N | F | P | P | D | G | P | V | M | Q | K | K | T | M | G | W | E | A | S | T | E | R |
| **BRIPO0.5** | Y | K | V | K | L | R | G | T | N | F | P | P | D | G | P | V | M | Q | K | K | T | M | G | W | E | A | S | T | E | R |
| **BRIPO** | Y | K | V | K | L | R | G | T | N | F | P | P | D | G | P | V | M | Q | K | K | T | M | G | W | E | A | S | T | E | R |

| | 151 | 152 | 153 | 154 | 155 | 156 | 157 | 158 | 159 | 160 | 161 | 162 | 163 | 164 | 165 | 166 | 167 | 168 | 169 | 170 | 171 | 172 | 173 | 174 | 175 | 176 | 177 | 178 | 179 | 180 |
|---|---|---|---|---|---|---|---|---|---|---|---|---|---|---|---|---|---|---|---|---|---|---|---|---|---|---|---|---|---|---|
| **BREP** | L | Y | P | E | D | G | V | L | K | G | D | I | K | M | A | L | R | L | K | D | G | G | R | Y | L | A | D | F | K | T |
| **BRIPO0.5** | L | Y | P | E | D | G | V | L | K | G | D | I | K | M | A | L | R | L | K | D | G | G | R | Y | L | A | D | F | K | T |
| **BRIPO** | L | Y | P | E | D | G | V | L | K | G | D | I | K | M | A | L | R | L | K | D | G | G | R | Y | L | A | D | F | K | T |

| | 181 | 182 | 183 | 184 | 185 | 186 | 187 | 188 | 189 | 190 | 191 | 192 | 193 | 194 | 195 | 196 | 197 | 198 | 199 | 200 | 201 | 202 | 203 | 204 | 205 | 206 | 207 | 208 | 209 | 210 |
|---|---|---|---|---|---|---|---|---|---|---|---|---|---|---|---|---|---|---|---|---|---|---|---|---|---|---|---|---|---|---|
| **BREP** | T | Y | K | A | K | K | P | V | Q | M | P | G | A | Y | N | V | D | R | K | L | D | I | T | S | H | N | E | D | Y | T |
| **BRIPO0.5** | T | Y | K | A | K | K | P | V | Q | M | P | G | A | Y | N | V | D | R | K | L | D | I | T | S | H | N | E | D | Y | T |
| **BRIPO** | T | Y | K | A | K | K | P | V | Q | M | P | G | A | Y | N | V | D | R | K | L | D | I | T | S | H | N | E | D | Y | T |

| | 211 | 212 | 213 | 214 | 215 | 216 | 217 | 218 | 219 | 220 | 221 | 222 | 223 | 224 | 225 | 226 | 227 | 228 | 229 | 230 | 231 | 232 | 233 | 234 | 235 | 236 | 237 | 238 | 239 | 240 |
|---|---|---|---|---|---|---|---|---|---|---|---|---|---|---|---|---|---|---|---|---|---|---|---|---|---|---|---|---|---|---|
| **BREP** | V | V | E | Q | Y | E | R | S | E | G | R | H | L | D | T | L | E | D | F | V | G | D | W | R | Q | T | A | G | Y | N |
| **BRIPO0.5** | V | V | E | Q | Y | E | R | S | E | G | R | H | L | V | T | L | E | D | F | V | G | D | W | R | Q | T | A | G | Y | N |
| **BRIPO** | V | V | E | Q | Y | E | R | S | E | G | R | H | L | V | T | L | E | D | F | V | G | D | W | R | Q | T | A | G | Y | N |

| | 241 | 242 | 243 | 244 | 245 | 246 | 247 | 248 | 249 | 250 | 251 | 252 | 253 | 254 | 255 | 256 | 257 | 258 | 259 | 260 | 261 | 262 | 263 | 264 | 265 | 266 | 267 | 268 | 269 | 270 |
|---|---|---|---|---|---|---|---|---|---|---|---|---|---|---|---|---|---|---|---|---|---|---|---|---|---|---|---|---|---|---|
| **BREP** | L | S | Q | V | L | E | Q | G | G | V | S | S | L | F | Q | N | L | G | V | S | V | T | P | I | Q | R | I | V | L | S |
| **BRIPO0.5** | L | S | Q | V | L | E | Q | G | G | V | S | S | L | F | Q | N | L | G | V | S | V | T | P | I | Q | R | I | V | L | S |
| **BRIPO** | L | S | Q | V | L | E | Q | G | G | V | S | S | L | F | Q | N | L | G | V | R | W | T | P | I | Q | R | I | V | L | S |

| | 271 | 272 | 273 | 274 | 275 | 276 | 277 | 278 | 279 | 280 | 281 | 282 | 283 | 284 | 285 | 286 | 287 | 288 | 289 | 290 | 291 | 292 | 293 | 294 | 295 | 296 | 297 | 298 | 299 | 300 |
|---|---|---|---|---|---|---|---|---|---|---|---|---|---|---|---|---|---|---|---|---|---|---|---|---|---|---|---|---|---|---|
| **BREP** | G | E | N | G | L | K | I | D | I | H | V | I | I | P | Y | E | G | L | S | G | D | Q | M | G | Q | I | E | K | I | F |
| **BRIPO0.5** | G | G | N | G | L | K | I | G | I | H | V | I | I | P | Y | E | G | L | S | G | D | Q | M | G | Q | I | E | K | I | F |
| **BRIPO** | G | G | N | G | L | K | I | G | I | H | V | I | I | P | Y | E | G | L | S | G | D | Q | M | G | Q | I | E | K | I | F |

| | 301 | 302 | 303 | 304 | 305 | 306 | 307 | 308 | 309 | 310 | 311 | 312 | 313 | 314 | 315 | 316 | 317 | 318 | 319 | 320 | 321 | 322 | 323 | 324 | 325 | 326 | 327 | 328 | 329 | 330 |
|---|---|---|---|---|---|---|---|---|---|---|---|---|---|---|---|---|---|---|---|---|---|---|---|---|---|---|---|---|---|---|
| **BREP** | K | V | V | Y | P | V | D | N | H | H | F | K | V | I | L | H | Y | G | T | L | V | I | D | G | V | T | P | N | M | I |
| **BRIPO0.5** | K | V | V | Y | P | V | D | N | H | H | F | K | V | I | L | H | Y | G | T | L | V | I | D | G | V | T | P | N | M | I |
| **BRIPO** | K | V | V | Y | P | V | D | N | H | H | F | K | V | I | L | H | Y | G | T | L | V | I | D | G | V | T | P | N | M | I |

| | 331 | 332 | 333 | 334 | 335 | 336 | 337 | 338 | 339 | 340 | 341 | 342 | 343 | 344 | 345 | 346 | 347 | 348 | 349 | 350 | 351 | 352 | 353 | 354 | 355 | 356 | 357 | 358 | 359 | 360 |
|---|---|---|---|---|---|---|---|---|---|---|---|---|---|---|---|---|---|---|---|---|---|---|---|---|---|---|---|---|---|---|
| **BREP** | D | Y | F | G | R | P | Y | E | G | I | A | V | F | D | G | K | K | I | T | V | T | G | T | L | W | N | G | N | K | I |
| **BRIPO0.5** | D | Y | F | G | R | P | Y | E | G | I | A | V | F | D | G | K | K | I | T | V | T | G | T | L | W | N | G | N | K | I |
| **BRIPO** | D | Y | F | G | R | P | Y | E | G | I | A | V | F | D | G | K | K | I | T | V | T | G | T | L | W | N | G | N | K | I |

| | 361 | 362 | 363 | 364 | 365 | 366 | 367 | 368 | 369 | 370 | 371 | 372 | 373 | 374 | 375 | 376 | 377 | 378 | 379 | 380 | 381 | 382 | 383 | 384 | 385 | 386 | 387 | 388 | 389 | 390 |
|---|---|---|---|---|---|---|---|---|---|---|---|---|---|---|---|---|---|---|---|---|---|---|---|---|---|---|---|---|---|---|
| **BREP** | I | D | E | R | L | I | N | P | D | G | S | L | L | F | R | V | T | I | N | G | V | T | G | W | R | L | H | E | R | I |
| **BRIPO0.5** | I | D | E | R | L | I | N | P | D | G | S | L | L | F | R | V | T | I | N | G | V | T | G | W | R | L | H | E | R | I |
| **BRIPO** | I | D | E | R | L | I | N | P | D | G | S | L | L | F | R | V | T | I | N | G | V | T | G | W | R | L | H | E | R | I |

| | 391 | 392 |
|---|---|---|
| **BREP** | L | A |
| **BRIPO0.5** | L | A |
| **BRIPO** | L | A |

**Figure S3. Sequence alignment of BREP, BRIPO0.5 and BRIPO.** Sequences derived from mScarlet-I and teLuc are colored in red and cyan, respectively. Linker residues are colored in gray. Mutations gained during protein engineering are shaded in yellow (from random mutagenesis) and orange (from site-directed randomization).

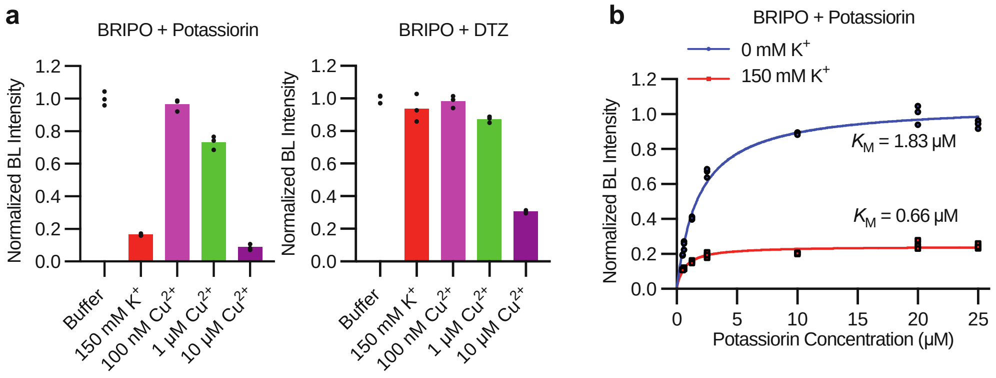


**Figure S4. Additional *in vitro* characterization of BRIPO.** (**a**) Bioluminescence responses of BRIPO in the presence of potassiorin (left) or DTZ (right) to 150 mM $K^+$ and the indicated concentrations of $Cu^{2+}$. n = 3 technical repeats. (**b**) Potassiorin concentration dependency of BRIPO bioluminescence at 590 nm in the presence or absence of 150 mM KCl. n = 3 technical repeats. The Michaelis-Menten equation was used to fit the data and derive the Michaelis constants ($K_M$). BL, bioluminescence.

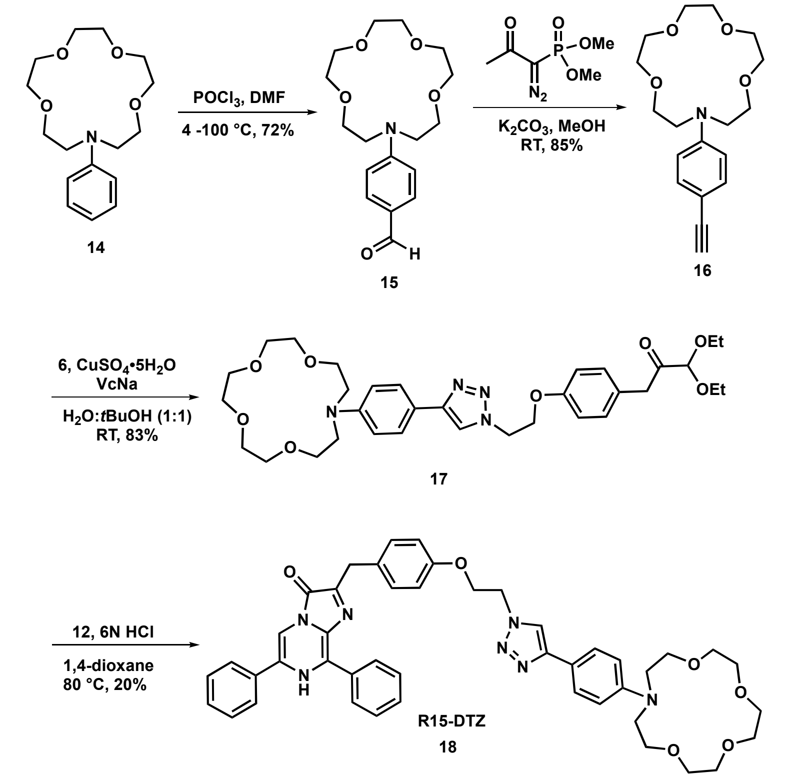


**Figure S5. Synthetic route to chemically prepare R15-DTZ.**

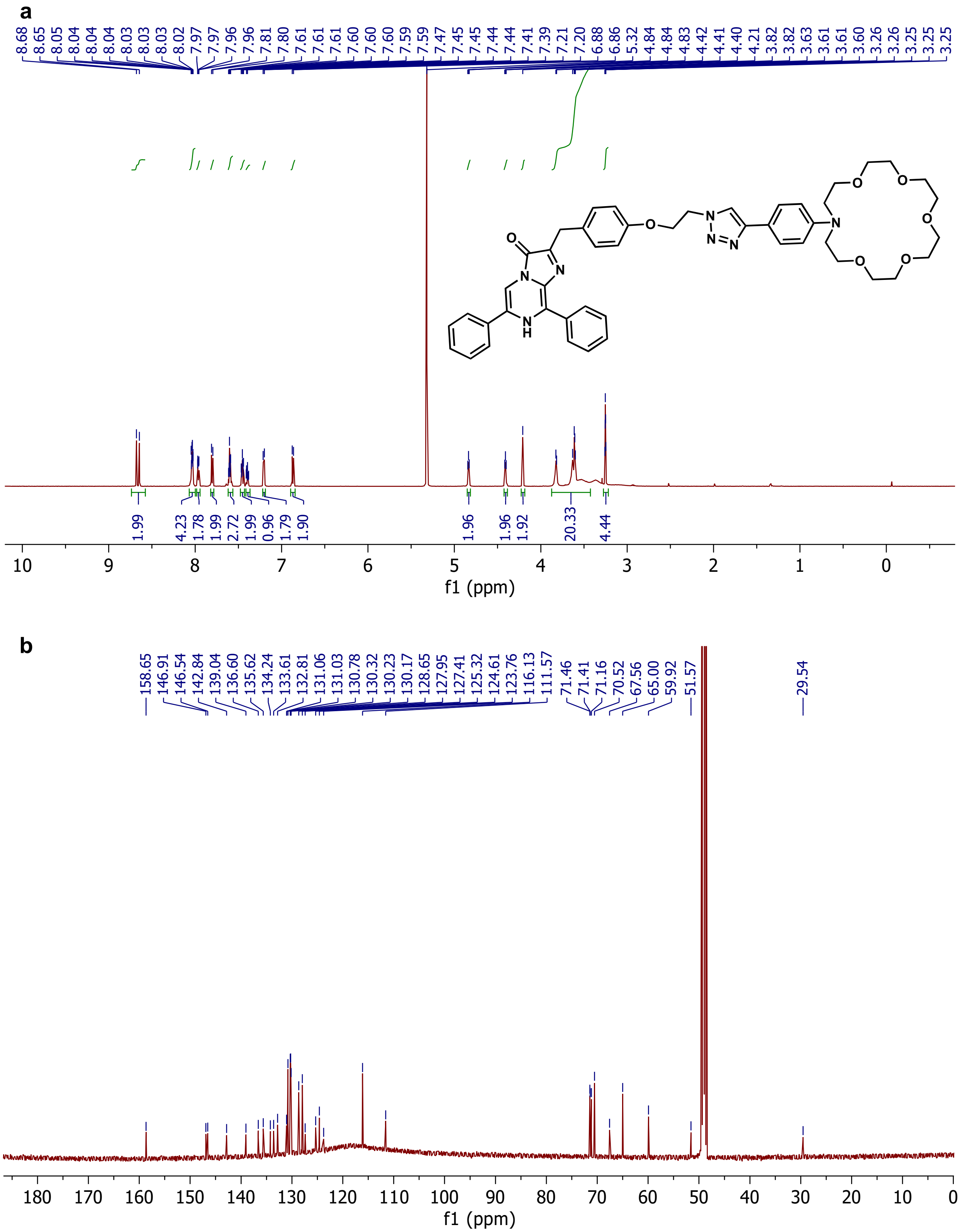


**Figure S6.** **$^{1}$H-NMR (a) and $^{13}$C-NMR (b) spectra for compound 11.**

a

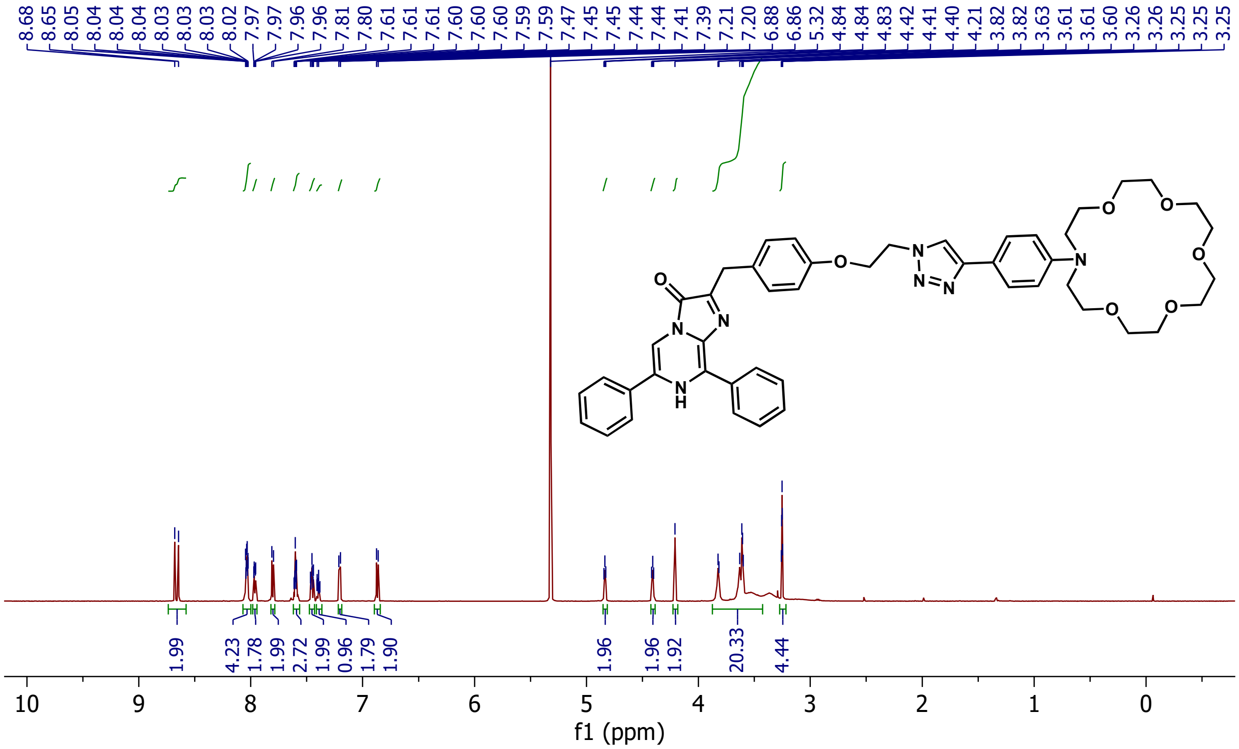

b

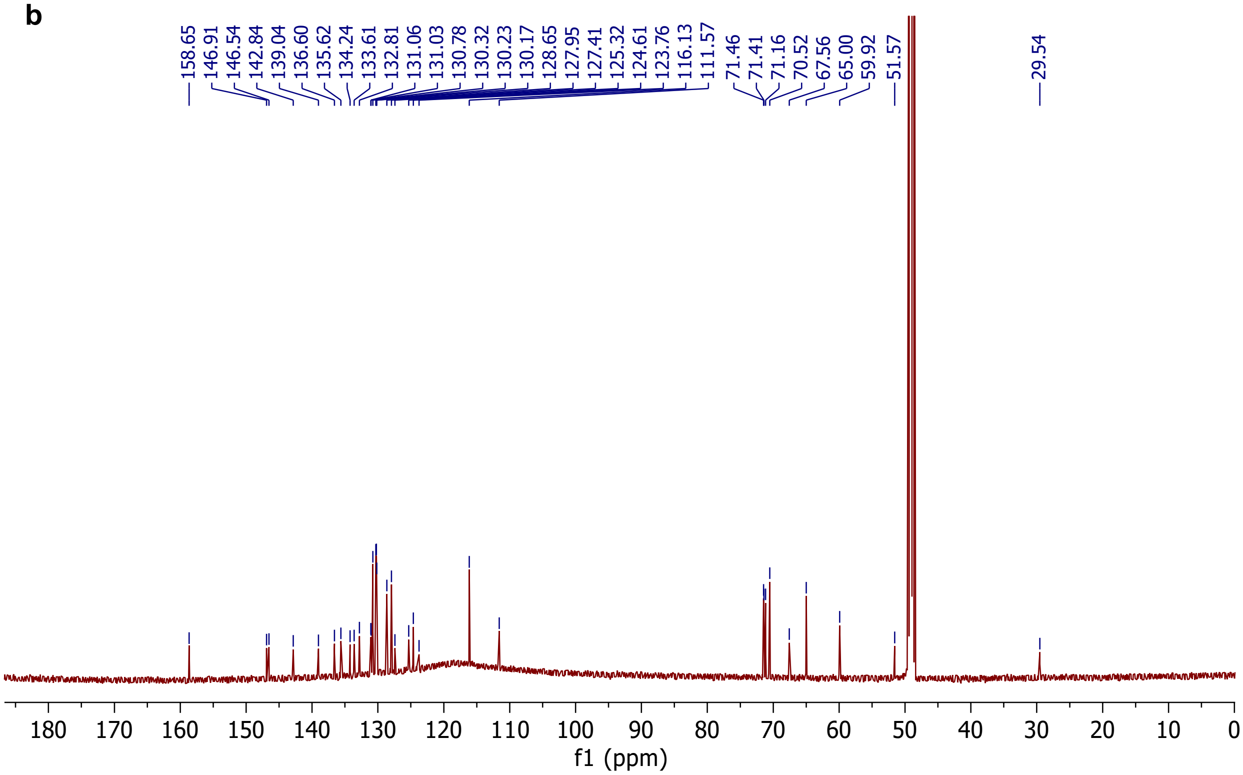

**Figure S7. $^{1}$H-NMR (a) and $^{13}$C-NMR (b) spectra for potassiorin (compound 13).**

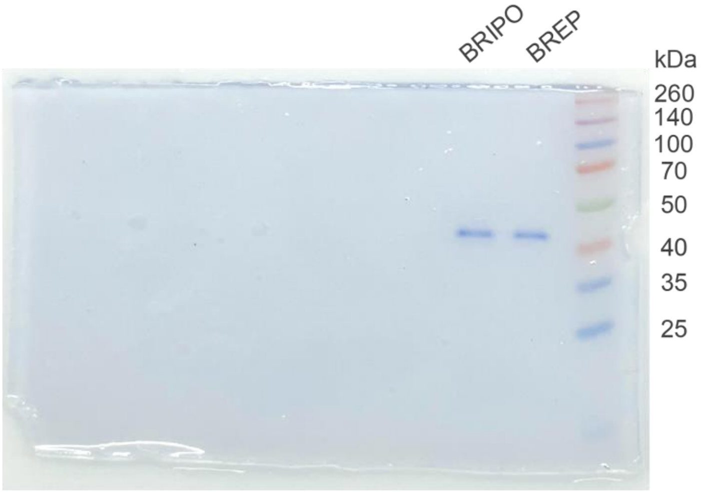


**Figure S8. SDS-PAGE showing the purity of the prepared BREP and BRIPO proteins.**

**Movie S1.**
Description: BLI of BRIPO-expressing HEK 293T cells in response to a combination of nigericin, ouabain, and bumetanide.

**Movie S2.**
Description: BLI of BRIPO-expressing HEK 293T cells stably expressing mTrek and several other ion channels in response to arachidonic acid.

**Movie S3.**
Description: BLI of a mouse with BRIPO expression in the brain in response to local glutamate stimulation.
